# Optogenetic manipulation of cGMP highlights PDE5 as the predominant cGMP-hydrolyzing PDE in megakaryocytes

**DOI:** 10.1101/2021.11.12.468375

**Authors:** Yujing Zhang, Pascal Benz, Daniel Stehle, Shang Yang, Hendrikje Kurz, Susanne Feil, Georg Nagel, Robert Feil, Shiqiang Gao, Markus Bender

## Abstract

Cyclic guanosine monophosphate (cGMP) signalling plays a fundamental role in many cell types including platelets. cGMP has been implicated in platelet formation, but mechanistic detail about its spatiotemporal regulation in megakaryocytes (MKs) is lacking. We expressed a photo-activated guanylyl cyclase, *Blastocladiella emersonii* Cyclase opsin (*Be*Cyclop), after viral-mediated gene transfer in bone marrow (BM)-derived MKs to precisely light-modulate cGMP levels. *Be*Cyclop-MKs showed a significantly increased cGMP concentration after illumination, which was strongly dependent on phosphodiesterase (PDE) 5 activity. This finding was corroborated by real-time imaging of cGMP signals which revealed that pharmacological PDE5 inhibition also potentiated nitric oxide (NO) triggered cGMP generation in BM MKs. In summary, we established for the first time optogenetics in primary MKs and identified PDE5 as the predominant PDE regulating cGMP levels in MKs. These findings also demonstrate that optogenetics allows for the precise manipulation of MK biology.

## Introduction

Binding of nitric oxide (NO) to guanylyl cyclases increases the cGMP formation. cGMP is a key intracellular signalling molecule in many cell types and tissues and exerts multiple cellular effects via its downstream effectors, such as protein kinase G (PKG) or cyclic nucleotide-gated (CNG) channels (1). The cGMP signalling pathway is long known for its critical role in the maintenance of cardiovascular homeostasis. Fluorescent cGMP biosensors have emerged as powerful tools for the sensitive analysis of cGMP pathways at the single-cell level (2), and methods that allow for precise manipulation of cGMP levels by light are being developed (3). The NO/cGMP/PKG pathway is highly expressed in platelets and its activation has been linked to platelet inhibition (4). Analysis of spatiotemporal dynamics of platelet cGMP using cGMP sensor mice revealed that cGMP generation is increased in shear-exposed platelets at the periphery of a thrombus thereby limiting thrombus growth (5). Cellular cGMP levels are regulated by phosphodiesterases (PDEs), which catalyse the hydrolysis of the 3′ phosphate bond of cGMP to generate 5′ GMP. Platelets express PDE2, 3 and 5, of which each is able to degrade cGMP (4-7). Bone marrow (BM) megakaryocytes (MKs) are the immediate progenitor cells of blood platelets (8). Studies on MKs derived from mouse fetal liver cells showed that PDE3A, PDE4A1 and PDE5 are detectable in maturing MKs, while PDE2A is found in the non-MK fraction, and that *in vitro* platelet release is enhanced by cGMP (9). However, the spatiotemporal regulation of cGMP signals by PDEs in MKs has not been addressed due to the lack of appropriate tools. Therefore, we expressed a light-sensitive guanylyl cyclase in primary BM-derived MKs to tightly manipulate cGMP levels (3). We demonstrate that the cGMP concentration in these MKs can be increased upon illumination, which rapidly declines in the dark caused by PDE5 activity.

## Results and Discussion

Concentration-dependent increase of cGMP in BM-derived MKs was measured when cells had been incubated with riociguat, a pharmacological stimulator of the NO-sensitive guanylyl cyclase (Fig. 1A). To tightly manipulate cGMP levels, the photoactivated guanylyl cyclase opsin from *Blastocladiella emersonii, Be*Cyclop (Fig. 1B), was expressed in BM-derived MKs after viral transduction, which allows light-triggered cGMP increase as shown in heterologous cells and *Caenorhabditis elegans* (3). Localization of YFP-*Be*Cyclop in the membrane system of BM-derived MKs was confirmed by confocal microscopy (Fig. 1C). Illumination (green light: 520 nm, 5 min) of YFP-*Be*Cyclop MKs resulted in a significant increase in cGMP (3.25±0.43 pmol/mL) as compared to samples in the dark (0.76±0.06 pmol/mL) and untransduced cells (no light: 0.95±0.09 pmol/mL; light: 0.84±0.02 pmol/mL) (Fig. 1D). These data demonstrate that it is possible to modulate intracellular cGMP in primary MKs with optogenetic approaches. Stimulation with 5 µM riociguat resulted in a similar cGMP increase as observed after 5 min illumination of YFP-*Be*Cyclop MKs (Fig. 1D). However, it has to be noted that direct comparison cannot be made since the transduction efficiency of YFP-*Be*Cyclop was only about ∼30% in MKs suggesting even higher cGMP values per cell after illumination. Interestingly, a rapid decrease of the intracellular cGMP concentration close to the baseline level was observed within 3 min after illumination, pointing to a high cGMP-hydrolyzing PDE activity in MKs (Fig. 1E). Copy number analysis of the murine platelet proteome revealed a high PDE5 expression in murine platelets (10). Therefore, cells were pretreated with the specific PDE5 inhibitor tadalafil, which resulted in a significant but only subtle cGMP increase (DMSO: 1.02±0.15 pmol/mL; tadalafil: 1.34±0.09 pmol/mL; Fig. 1F). However, cGMP levels were 6.9-fold increased when pretreated with tadalafil and 5 min illuminated (20.45±4.3 pmol/mL), as compared to illumination without tadalafil pretreatment (2.95±0.45 pmol/mL). These data strongly suggest that PDE5 limits the cGMP concentration increase in MKs. Furthermore, inhibition of PDE5 resulted in less cGMP decrease to 10±1.5 pmol/mL in 3 min after illumination (Fig. 1F), again comfirming that PDE5 degrades cGMP in MKs. In agreement with our findings with tadalafil, illuminated MKs preincubated with other PDE5 inhibitors (vardenafil: PDE5, 6; sildenafil: PDE5, 6) also significantly increased cGMP concentration compared to illuminated YFP-*Be*Cyclop MKs without inhibitor (Fig. 1G,H), whereas non-PDE5 inhibitors did not elevate intracellular cGMP (vinpocetine: PDE1; EHNA: PDE2; BAY60-7550: PDE2; milrinone: PDE3, 4; TAK-063: PDE10; Fig. 1G). Interestingly, also the broad-spectrum PDE inhibitors zaprinast (PDE5, 6, 9, 10, 11) and IBMX (PDE1, 2, 3, 4, 5, 7, 11) had no or only minor effects, respectively (Fig. 1G).

**Figure 1.**
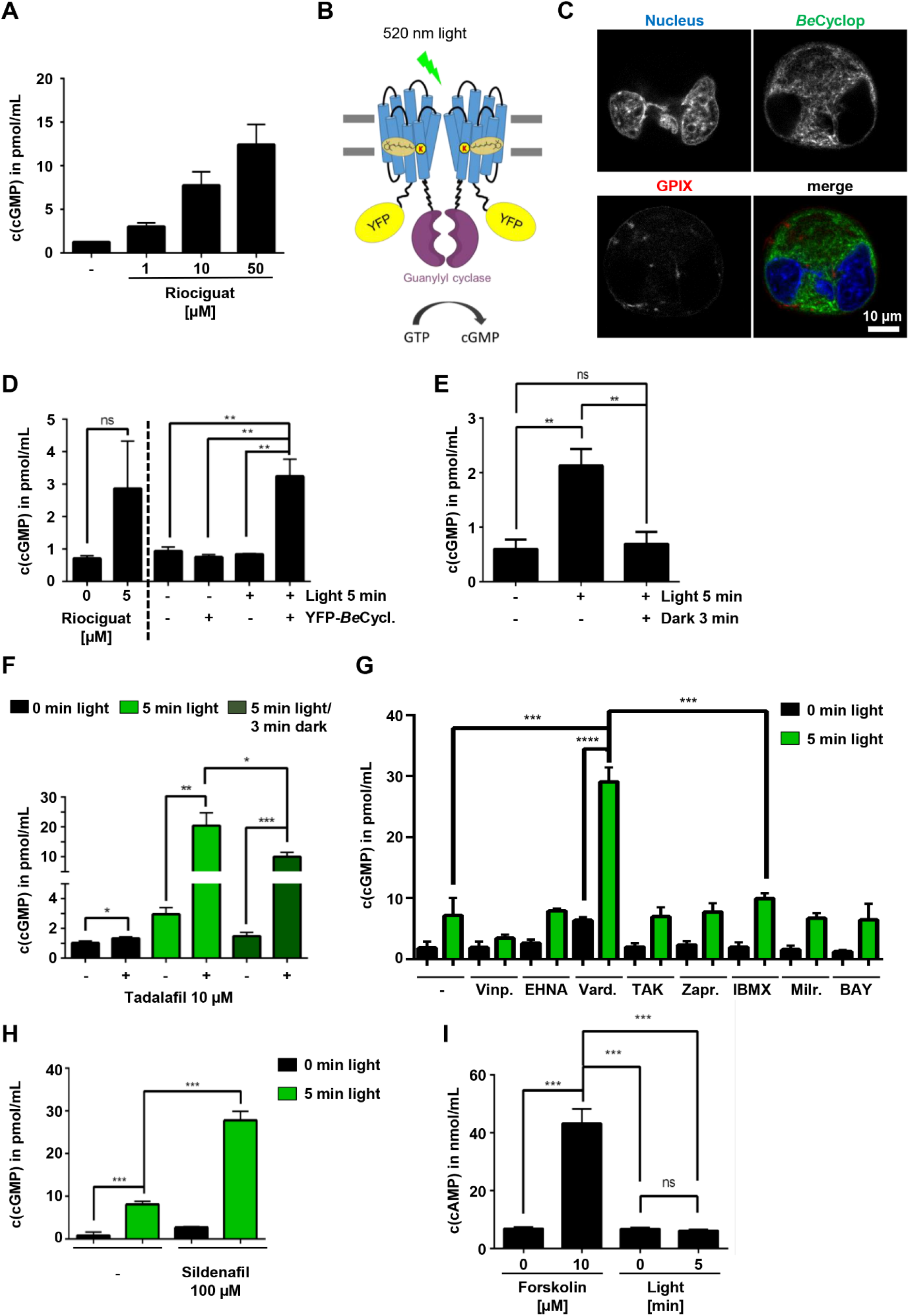
PDE5 regulates optogenetically-increased cGMP levels in MKs. (A) BM-derived MKs were incubated for 10 min with 1, 10, or 50 μM riociguat. DMSO was used as vehicle control. (n=3). (B) A schematic model of the light-gated guanylyl cyclase, *Be*Cyclop. (C) Representative images (microscope: Leica SP8) of YFP-*Be*Cyclop expression in BM-derived MKs. MK/platelet-specific membrane protein glycoprotein (GP) IX. (D) BM-derived YFP-*Be*Cyclop-expressing MKs were illuminated with green light (520 nm) for 5 min and immediately lysed. Untransduced cells or no illumination served as controls. (n=3, representative for 2 independent experiments). (E) YFP-*Be*Cyclop-expressing MKs were illuminated with green light (520 nm) for 5 min and immediately lysed or after 3 min in the dark. (F) BM-derived MKs were preincubated with tadalafil for 10 min prior to 5 min illumination. Cells were either lysed immediately or after 3 min before measuring cGMP concentration. (n=3, representative for 2 independent experiments). (G, H) BM-derived MKs were preincubated with different PDE inhibitors for 10 min prior to illumination. Cells were either lysed immediately or after 3 min in the dark before measuring cGMP concentration. DMSO control in (G), H_2_0 control in (H). (n=3). Vinpocetine (Vinp.): 20 µM, EHNA: 10 µM, vardenafil (Vard.): 50 µM, TAK-063: 0.15 µM, zaprinast (Zapr.): 40 µM, IBMX: 100 µM, milrinone (Milr.): 20 µM, BAY 60-7550: 10 µM, sildenafil: 100 µM. (I) Determination of cAMP concentration of YFP-*Be*Cyclop-expressing BM-derived MKs after incubation with forskolin or illumination with green light for 5 min (n=3).

To demonstrate the high specificity of *Be*Cyclop, MKs expressing YFP-*Be*Cyclop were either stimulated with the adenylyl cyclase activator forskolin or illuminated and subsequently, cyclic adenosine monophosphate (cAMP) was determined. While forskolin significantly increased the cAMP concentration in MKs, illumination had no effect on cAMP concentration, demonstrating that *Be*Cyclop cannot produce or influence cAMP (Fig. 1I).

Next, we used MK/platelet-specific cGMP sensor mice to spatiotemporally visualize cGMP dynamics in MKs *ex vivo* (Fig. 2A) (5, 11). Isolated femurs with externalized bone marrow were superfused with 1 mL/min imaging buffer. In contrast to static cell culture experiments, this method allows to rapidly add and remove the pharmacological compounds within seconds similar to optogenetic experiments. Application of the NO donor diethylamine NONOate (DEA/NO) increased cGMP in BM MKs (Fig. 2B,C). Similar to the optogenetic results, the PDE5-specific inhibitor tadalafil alone slowly increased cGMP and significantly augmented the NO-induced cGMP signal in MKs (Fig. 2B,C). These data strongly suggest that BM MKs in their native environment express a functional NO/cGMP/PDE5 signalling pathway that can be pharmacologically enhanced with PDE5 inhibitors.

**Figure 2.**
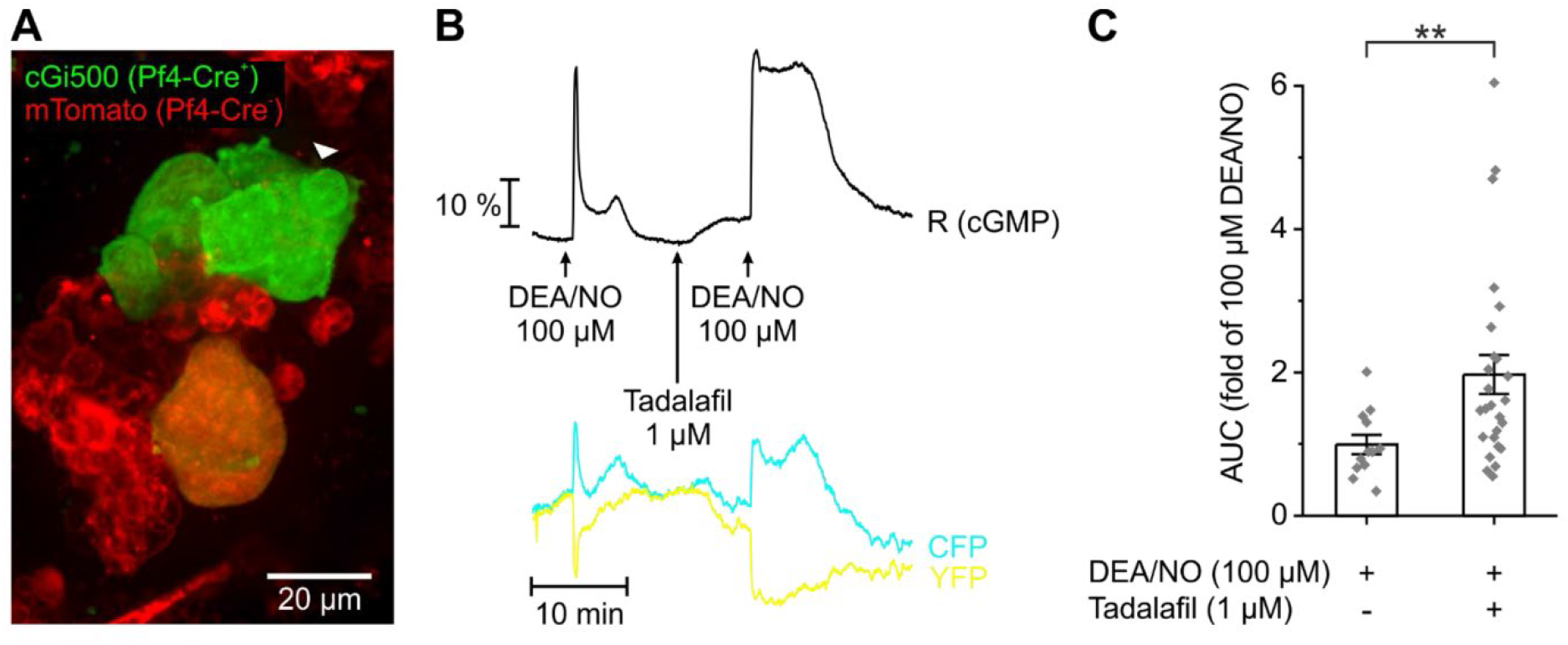
Real-time cGMP imaging in MKs. FRET-based cGMP imaging was performed with BM from MK/platelet-specific cGMP sensor mice. (A) Representative field of view of a measurement. Green color indicates cGMP sensor expressing MKs. mTomato (red) is expressed in all other cells. The white arrowhead points towards the MK measured in (B). (B) Representative real-time FRET/cGMP measurement. During recording, DEA/NO (100 µM) was applied for 2 min with or without 7 min pre-incubation with tadalafil (1 µM). Black trace represents the CFP/YFP ratio R, which indicates cGMP concentration changes. Relative fluorescence changes of the individual fluorescent proteins are shown as cyan (CFP) and yellow (YFP) trace. (C) Statistical analysis was performed with the area under the curve (AUC) of cGMP signals induced by DEA/NO (100 µM) in the presence of tadalafil (1 µM) or vehicle (DMSO; 0.002%). Data represent mean ± SEM of n ≥ 12 MKs from ≥ 3 mice. Scale bar, 20 µm.

In summary, we established to the best of our knowledge the first time optogenetics in primary MKs. By expressing the photo-activated guanylyl cyclase, *Be*Cyclop, we tightly controlled cGMP levels in MKs by light. We identified PDE5 as the major PDE, counteracting cGMP levels in BM-derived MKs, and could verify our optogenetic results by cGMP imaging in BM MKs *ex vivo* using cGMP sensor mice. Our findings are corroborated by research on mouse fetal liver cell (FLC)-derived MKs showing PDE5 expression is highest in mature FLC-derived MKs (9). Our study shows that optogenetics in MKs allows to light-modulate the function of key signalling molecules thereby possibly identifying novel regulatory mechanisms of MK maturation and platelet production. However, our study has also limitations as the transduction efficiency of MKs was only about 30%. We analysed the complete MK fraction since complicated sorting of mature MKs (cell diameter of approx. 35 µm) most likely leads to light-activation before starting the experiment. Thus, our data underestimate the light-induced increase of cGMP in single MKs expressing YFP-*Be*Cyclop due to the untransduced cell fraction. Further studies are required using MK/platelet-specific optogenetic mouse lines to spatiotemporally light-manipulate molecules or proteins in MKs and platelets *in vivo*. The finding that BM MK express PDE5 also informs future therapeutic strategies to increase cGMP concentrations and potentially modulate platelet biogenesis with clinically used PDE5 inhibitors.

## Materials and Methods

### Animals

MK/platelet-specific cGMP sensor mice (*cGi500-L2*^*fl/fl*^; *Pf4-Cre*^*tg/+ (12)*^) were used to visualize cGMP signals in MKs in real-time (5, 11).

### Reagents

Riociguat (Selleck Chemicals), tadalafil (Sigma-Aldrich/Supelco), DMSO, forskolin (Sigma-Aldrich), vinpocetine, EHNA (Enzo Life Sciences), BAY60-7550, TAK-063 (Cayman Chemical), milrinone, vardenafil, zaprinast (Santa Cruz Biotechnology), sildenafil (Merck), IBMX (Thermo Fisher Scientific), DEA/NO (Axxora).

### BM-derived MKs

BM cells were obtained from femur and tibia of C57BL/6 mice by flushing, and lineage depletion was performed using an antibody cocktail of anti-mouse CD3, clone 17A2; anti-mouse Ly-6G/Ly-6C, clone RB6-8C5; anti-mouse CD11b, clone M1/70; anti-mouse CD45R/B220, clone RA3-6B2; anti-mouse TER-119/Erythroid cells, clone Ter-119 (1.5 μg of each antibody per mouse, Biolegend) and magnetic beads (Dynabeads® Untouched Mouse CD4 Cells, Invitrogen). Lineage-negative (Lin-) cells were cultured in DMEM medium (supplemented with 4 mM L-glutamine, 100 U mL-1 penicillin, 50 mg mL-1 streptomycin) containing 1% of recombinant TPO (homemade) and 100 U mL-1 recombinant Hirudin (Hyphen Biomed) at 37°C under 5% CO_2_.

### Expression of YFP-*Be*Cyclop in MKs

YFP-*Be*Cyclop DNA was cloned into the murine stem cell virus (MSCV) vector and transfected into 293T cells. Viral supernatant was collected and BM-derived cells were infected on day 1 (13). A bovine serum albumin density gradient was used at culture day 3 to separate MKs from non-MK cells. Experiments were performed on day 4.

### Illumination and determination of cGMP concentration

Cell density was adjusted to 5 × 10^4^ cells/mL. If needed, samples were preincubated with agonist or inhibitor for 10 min at room temperature. Subsequently, samples were kept in the dark or illuminated with light (520 nm, 20 µW/mm^2^) for 5 min at room temperature and lysed either immediately or after 3 min in the dark with 0.1 M hydrogen-chloride (HCl). Concentrations of cGMP or cAMP were measured by DetectX Direct cyclic GMP or AMP Enzyme Immunoassay Kit, respectively, and analysed by the optical density at 450 nm with a Thermo Scientific™ Multiskan microplate spectrophotometer.

### Real-time cGMP imaging in MKs *ex vivo*

Femurs and tibias were dissected from MK/platelet-specific cGMP sensor mice (*cGi500-L2*^*fl/fl*^; *Pf4-Cre*^*tg/+ (12)*^). Bone tissue was carefully removed with scissor and forceps on one side to give access to the bone marrow. Opened bones were placed in a superfusion chamber (RC 26, Warner Instruments), mounted with a Slice Hold-Down (SHD 26H/10, Warner Instruments) and continuously superfused with imaging buffer (140 mM NaCl, 5 mM KCl, 1.2 mM MgCl_2_, 2 mM CaCl_2_, 5 mM HEPES and 10 mM D-glucose; pH = 7.4) with or without added drugs. Real-time FRET/cGMP imaging was performed by recording CFP and YFP fluorescence as described in detail elsewhere (14). The relative CFP/YFP ratio change (black trace in the respective graph; referred to as R (cGMP)) correlates with the cGMP concentration change.

### Data analysis

Results are shown as mean ± SEM. Statistical significance was assessed by an unpaired t-test with GraphPad Prism software. P-values <0.05 were considered significant (^****^ <0.0001; ^***^ <0.001; ^**^ 0.001 to 0.01; ^*^ 0.01 to 0.05). ns: non-significant. Statistical analysis of cGMP sensor imaging data was performed with Origin 2019 (OriginLab, Northampton, MA, USA). Since data sets were not normally distributed, statistical differences were analysed non-parametrically by Mann-Whitney U Test.

## Data availability

All study data are in the article.

## Acknowledgments

The authors thank Sarah Seidel for the help with experiments. This work was supported by TR240 grant with project number 374031971 of the Deutsche Forschungsgemeinschaft (DFG; German Research Foundation).

